# The actin assembly regulator *toca-1* regulates collateral branching in *C. elegans*

**DOI:** 10.1101/2025.10.27.684948

**Authors:** Anna Brinck, Scott Dour, M. L. Nonet

## Abstract

Neuronal branching is essential for establishing complex neural circuits. Many axons and dendrites in the CNS branch, forming complex process arbors. Some of these branches are formed during initial outgrowth and often occur via bifurcation of the growth cones of the extending neuronal processes. However, others are formed *de novo* by branch extension from a previously existing neurite. This process is often referred to as collateral branching. Here, we investigate the molecular mechanisms underlying this process. We show that in *C. elegans*, collateral branching of the PLM neuron is mediated by an actin assembly process guided by <u>T</u>ransducer of <u>C</u>DC-42-<u>D</u>ependent <u>A</u>ctin assembly-1 (TOCA-1). This scaffolding protein is recruited to the branching site at the time of branch formation. *cdc-42* and the guanine nucleotide exchange factor *dock-11* are also required for branch formation. Biochemically, TOCA-1 has been demonstrated to recruit WASP-1 and activate the Arp2/3 complex to promote actin assembly. In vivo, both *wsp-1* and the Arp2/3 complex mutants also disrupt branch formation. While loss of TOCA-1 disrupts branching, it does not influence anterior-posterior (AP) or dorsal-ventral (DV) positioning of the branch, in contrast to previously defined branching regulators. Our data support a model in which TOCA-1 acts downstream of AP and DV positioning factors, directly orchestrating filopodial extension to form the nascent branch.

## Introduction

Precise neuronal connectivity is fundamental to nervous system function, and understanding the molecular mechanisms that control axon branching patterns represents a central challenge in developmental neurobiology. During development, axons not only extend primary projections that can bifurcate at the growth cones but also generate collateral branches that emerge *de novo* from the axon shaft, independent of the growth cone (Gallo, 2011; Armijo-Weingart and Gallo, 2017). For example, sensory neurons form collateral branches through the localized formation of actin-based filopodia along the axon shaft, which are subsequently invaded by microtubules to mature into stable branches (Gallo and Letourneau, 1998; Armijo-Weingart and Gallo, 2017). The establishment of appropriate neuronal circuits requires not only accurate axon guidance to target regions but also precise spatial control over where and when axons form collateral branches to diversify their connectivity patterns (Tessier-Lavigne and Goodman, 1996; Gallo, 2011; Kalil and Dent, 2014).

The nematode *Caenorhabditis elegans* has emerged as a useful model organism for studying neuronal development due to its simple nervous system comprising only 302 neurons with stereotyped morphologies and connections (White *et al*., 1986). The mechanosensory neurons, extensively characterized for their role in gentle touch sensation, exhibit highly stereotyped development that provides an excellent framework for investigating molecular mechanisms controlling neuronal connectivity (Chalfie and Sulston, 1981; Goodman, 2006).

Spatial precision in neuronal branching is exemplified by the PLM touch receptor neurons (TRNs) of *C. elegans*. The PLM neurons are a set of two bilaterally symmetric TRNs that mediate responses to gentle touch on the posterior half of the body (Chalfie and Sulston, 1981). Each consists of a cell body located in the tail ganglia that extends a short posterior neurite and a long anterior directed process (referred herein as the AP process) which maintains close association with the cuticle. The AP process extends to a stereotypic position just posterior to the vulva, defining a non-overlapping tiled sensory territory with the ALM neurons which mediate touch in the anterior half of the animal (Gallegos and Bargmann, 2004; Chen *et al*., 2017). The AP process is filled with specialized 15-protofilament microtubules essential for mechanosensation, with mechanoreceptor complexes localized along the entire length and linked extracellularly to the collagen cuticle matrix and intracellularly to the microtubule cytoskeleton (Chalfie and Thomson, 1979; Zhang *et al*., 2002; Emtage *et al*., 2004). Activation of PLM neurons induces behavioral responses mediated by a combination of gap junction and chemical synaptic connections to command neurons that regulate locomotion. Remarkably, these connection types exhibit precise spatial segregation. Gap junction connections are located on the AP process just anterior of the soma, while chemical synapses are restricted to a single synaptic branch that extends ventrally from the AP process at the stereotyped position approximately two-thirds of the distance between the soma and process ending, about 30-70 μm posterior to the vulva (Chalfie *et al*., 1985; Chen *et al*., 2017).

The development of PLM connectivity follows a precise temporal program. The AP process extends late in embryogenesis and its position relative to the ALM soma is established in the first few hours hatching and then grows interstitially as the animal develops to adulthood (Chalfie and Sulston, 1981; Gallegos and Bargmann, 2004). The synaptic branch develops from an initial filapodial protrusion that appears in the first few hours after hatching and subsequently extends ventrally and enters the ventral cord where it contacts command interneurons and forms a stable GFP-rab-3 containing varicosity by 12 hours post-hatching (Chalfie *et al*., 1985; Chen *et al*., 2014; Luo *et al*., 2014). The varicosity eventually grows to 5-7 microns in length by the late L4 stage (Schaefer *et al*., 2000). Each PLM extends only a single branch, demonstrating remarkable spatial precision in branch positioning. This stereotyped branching pattern, with both the AP process terminus and synaptic branch occurring at reproducible positions, exemplifies the precision of developmental programs that establish neuronal connectivity (Chen *et al*., 2017).

This spatial precision in PLM branching reflects broader principles of collateral branch formation, a key mechanism by which neurons diversify their output and establish complex connectivity patterns. Unlike primary axonal outgrowth guided by growth cones during early development, collateral branches emerge from post-embryonic axons through distinct processes involving cytoskeletal remodeling (Gibson and Ma, 2011; Kalil and Dent, 2014). Guidance molecules such as netrin are known to influence branching decisions (Dent *et al*., 2004; Hao *et al*., 2010; Chen *et al*., 2017), but the challenge lies in understanding how these spatial cues are translated into localized cellular responses with the precision required for stereotyped connectivity.

Axonal branch formation requires precise coordination of both actin and microtubule dynamics. Current models propose that collateral branching begins when F-actin patches generate filopodia, which are subsequently stabilized by microtubule invasion (Chia *et al*., 2012; Armijo-Weingart and Gallo, 2017; Chen *et al*., 2017). One mechanism depends on Wiskott-Aldrich syndrome protein (WASP) and the Arp2/3 complex to create branched actin networks at membrane protrusion sites with WASP activation controlled by the small GTPase CDC-42 (Millard *et al*., 2004; Rotty *et al*., 2013).

A critical gap in our understanding is how these general cytoskeletal mechanisms achieve spatial precision in PLM neurons. The reproducible positioning of the PLM branch indicates that spatially encoded signals must direct where branches form, yet how molecular components are organized in space and time remains unclear.

To address this question, we used genetic analysis to identify crucial components of PLM branching. Through this approach, we identified TOCA-1, a member of the highly conserved PCH (*pombe* Cdc15 homology) protein family, that consists of F-BAR, coiled-coil, and SH3 domains that enable it to simultaneously bind membrane phospholipids, recruit CDC-42, and activate WASP (Ho *et al*., 2004; Bu *et al*., 2009; Fricke *et al*., 2009; Giuliani *et al*., 2009). We hypothesize that TOCA-1 functions as a spatial organizer: its F-BAR domain marks future branch sites through membrane interactions, while its coiled-coil and SH3 domains coordinate CDC-42 and WASP recruitment to ensure branches form at precise axonal positions.

Using live imaging and epistatic analysis, we investigated TOCA-1’s role in PLM branching. Our findings demonstrate that TOCA-1 functions cell-autonomously in PLM neurons and is essential for collateral branch formation without affecting other types of neuronal branching. TOCA-1 acts with the putative CDC-42 Guanine nucleotide Exchange Factor DOCK-11, CDC-42, WSP-1 (the *C. elegans* WASP homolog), and the Arp2/3 complex in a coordinated pathway that integrates with netrin signaling and membrane trafficking to initiate branching at the correct spatial location. These results support a model in which TOCA-1 serves as a central molecular scaffold that coordinates multiple regulatory inputs to achieve the spatial precision required for stereotyped neuronal connectivity patterns.

## Materials and Methods

### *C. elegans* strain maintenance

*C. elegans* was grown on NGM plates seeded with *E. coli* OP-50 on 6-cm plates. Stock strains were maintained at room temperature (∼22.5°C). The *js205* allele was isolated in a clonal screen as described in Nonet (1999) and (Schaefer, 2001) Strains used in this study are listed in Table S1. Additionally, *F22G12.5* was renamed to *dock-11* following approval from the *C. elegans* gene naming committee, based on its orthology to human DOCK11 and the presence of characteristic DOCK family domains including the N-terminal PH domain that distinguishes it from other DOCK family members

### Plasmid constructions

Golden Gate reactions were performed as described in Nonet (2020). In-Fusion reactions were performed according to manufacturer’s protocols (Takara Bio). SapTrap assembly was performed as described in Knoebel *et al*. (2023). A detailed description of individual constructions is provided in Supplemental Methods. DNA constructs and oligonucleotides used are listed in Table S1.

### CRISPR/cas9-mediated mutagenesis

CRISPR/cas9 genome editing was performed using methods adapted from Arribere *et al*. (2014). sgRNA expression plasmids were constructed by ligating annealed oligonucleotides into BsaI-digested pRB1017 (Addgene #59936). Cas9 expression was driven by peft-3-cas9 (NMp3143). DNA mixtures were injected at 50 ng/μl of the Cas9 plasmid, 20 ng/μl of the dyp-10 sgRNA (NMp3153), 500nM of the dpy-10 repair oligo (NMo5238), 37.5-50 ng/μl of the sgRNA target, and 500nM of the target repair oligo, where applicable. Injected animals were screened for desired mutations using PCR amplification and restriction analysis then sequencing. All mutant alleles created for this study are listed in Table S1.

### Generation of transgenes

Microinjections were performed as described in Nonet (2020). A paint brush (Robert Simmons E51 liner 10/0) was used to mount animals onto agar pads before injection. Animals were typically injected in a single gonad. DNA were injected at ∼50 μg/ml in 10 mM Tris pH 8.0, 0.1 mM EDTA. miniMos insertions (Frøkjær-Jensen *et al*., 2014) were created as described in Nonet (2020). For RMCE integration (Nonet, 2023) and RMI (Beck and Nonet, 2025) injected P0 animals were pooled 2-3 per plate and maintained at 25°C. For Drug selection was applied 3 days after injection using hygromycin B (100 μl of 20 mg/ml) or G418 disulfate (500 μl of 25 mg/ml). Homozygous integrants were identified ∼ 8 days after injection. All transgenes created for this study are listed in Table S1.

### Microscopy and quantification

Screening of worms for fluorescence during analysis was performed on a Zeiss (White Plains, NY) Stemi SV11 dissecting microscope outfitted with a Lumencor (Beaverton, OR) Sola light source and a Kramer Scientific (Amesbury, MA) M2bio epi-fluorescence module with a long pass GFP filter and an RFP filter and a rotating turret housing both a Zeiss 1.6X objective and a Kramer 10X (n.a. 0.45) objective for high power observation. For imaging, animals were mounted with 1 mM levamisole and 25 micron polystyrene beads in PBS. 10-25 animals were typically placed on a single slide. Animals were imaged on an Olympus (Center Valley, PA) BX-60 microscope run using Micro-Manager 2.0ß software (Edelstein *et al*., 2014) using either a 20X air (n.a. 0.5) or 40X air (n.a. 0.75) plan-NEOFLUAR objective. The scope is equipped with a ToupTek MAX04BM sCMOS camera, a Lumencor AURA LED light source, an Applied Scientific Instruments (Eugene, OR) MS-4000 XY piezo Z motorized stage, Ludl (Hawthorne, NY) high speed electronic filter wheels and shutters and a Chroma 59022 eGFP/MCherry dual band filter set. Typically, at 20X, stacks of images at 2.5 um intervals were collected. Images were quantified using FIJI ImageJ software (Schindelin *et al*., 2012).

To quantify PLM branch number, only outgrowths contacting the ventral nerve cord were counted as branches in all animals except *mec-7* mutants. In *mec-7* null animals, substantial outgrowths that failed to contact the ventral cord were also counted as branches. For PVD branch density, maximum projection images were generated, and tertiary (4′) branches were counted along ∼100 μm of the dendrite. Branch density was calculated by dividing the number of branches by the length analyzed. To quantify *mito-*GFP and GFP-ELKS-1 in *toca-1* mutant animals (NM6847 and NM6848), animals were maintained at 25°C, and only unbranched PLM processes were selected. GFP-positive puncta were counted along ∼200 μm of the PLM process posterior to the vulva, and GFP density was calculated by dividing the number of puncta by the length of the process. For GFP-ELKS-1 fluorescence intensity analysis, ten puncta located posterior to the vulva were selected from each PLM process. The integrated density of these puncta was measured, normalized to background fluorescence, averaged per process, and plotted. For synapse intensity quantification, wild-type (NM3361) and *toca-1* mutant animals (NM6887) were maintained at 25°C. Maximum projection images were generated, and integrated density measurements were taken from equal areas within synaptic regions. Values were normalized to background fluorescence and plotted. To quantify synapse size, maximum intensity projections from wild-type (NM3361) and *toca-1* mutant (NM6887) animals maintained at 25°C were thresholded using identical parameters to highlight synaptic areas. Thresholded synapses were selected and their areas measured using the ‘Measure’ function and plotted. To measure PLM branch and ALM soma positions and account for body curvature, we constructed an anatomical landmark axis (see methods for details) using X and Y coordinates of anatomical landmarks: AVM soma (0%), vulva (50%), and PVM soma (100%). PLM branch and ALM soma positions were determined by projecting each structure onto this axis and calculating the percentage distance along that AP axis. Animals with fewer than two landmarks were excluded from analysis. Abnormal versus normal PVM and AVM morphology was quantified using a categorical scoring system, where abnormal phenotypes were defined as failure to meet the ventral nerve cord, absence of the neuron, incorrect process length, branching defects, migration paths with >45 degree turns, or ventral growth patterns (categories shown in Figure S8). GFP-RAB3 accumulations along mechanosensory processes were quantified using a binary scoring system, with accumulations designated as ‘present’ when 3 or more puncta were detected and ‘absent’ when less than 3 puncta were observed. All plots were generated using BioRender.

### RNAi feeding

RNAi feeding experiments were performed as described in Kamath *et al*. (2001). Bacterial clones expressing dsRNA were obtained from the Ahringer RNAi library (Kamath and Ahringer, 2003). HT115(DE3) bacteria were grown in LB + 50 μg/ml ampicillin for 6-12 hours, then seeded onto NGM plates containing 25 μg/ml carbenicillin and 1 mM IPTG. Plates were induced overnight at room temperature. L3-L4 hermaphrodites were transferred to feeding plates and maintained at 15°C for 72 hours before analysis.

### Auxin experiments

Auxin-inducible degradation experiments were performed using strains expressing AID-tagged proteins. For all strain carrying TIR1 except strain NM5031, L4s were placed on plates treated with 1 mM auxin (indole-3-acetic acid) dissolved in ethanol then L4 progeny were assayed. For strain NM5031, L4s were placed on plates treated with 100 μm of auxin. At this condition, progeny were assayed at L1 or L2 due to developmental arrest. For strains carrying TIR1(F79G), 1 μm 5-Ph-IAA (Sigma Millipore) was used. Control treatments used equivalent volumes of ethanol. For auxin time course experiments, animals were timed after hatching and placed on auxin treated plates at different intervals then assayed at L4 for branching phenotypes.

### Statistics

Statistical significance was denoted as ***p < 0.001, **p < 0.01, *p < 0.05, and n.s. = not significant. Asterisks located with the bars are comparing against wild-type animals. Branching pattern distributions were compared between genotypes using Mann-Whitney U tests with data detailed in Table S2. Due to unequal sample sizes between wild-type at 22°C (NM336, n=173) and mutant groups at 22°C, wild-type animals were randomly resampled without replacement to match mutant group sizes (n=50) to eliminate sample size bias. Mann-Whitney U tests were performed using these resampled wild-type counts against the original mutant data. Quantitative measures of *mec-7* aberrant branching, PVD branch density, *mec7p::GFP-elks-1*, *mec7p::mito-GFP*, GFP fluorescence and size of synapses, and PLM branch and ALM soma positions were analyzed using Welch’s t-tests to account for unequal variances. Chi-squared tests with Yates continuity were used to analyzed PVM and AVM phenotypes as well as to compare accumulations of RAB-3 on PLM processes. Error bars represent SEM in all figures except panels 7C-D. In these graphs, SD is used for error bars. Plots depict mean with SD.

## Results

PLMs are a bilateral pair of mechanosensory neurons that sense gentle touch in the posterior half of C. elegans animals. Each neuron extends a single anterior-directed process in a lateral position that is closely associated with the epidermis and terminates near the ALM soma, which senses touch in the anterior half of the animal. PLMs form both electrical and chemical synapses with postsynaptic partners. The chemical synapses are large multi-micron sized varicosities that form in the ventral nerve cord at the end of a single collateral branch extending ventrally from the antero-posterior sensory process at a stereotypic position approximately two-thirds of the distance from the soma to the process terminus (Fig. 1A).

**Figure 1.**
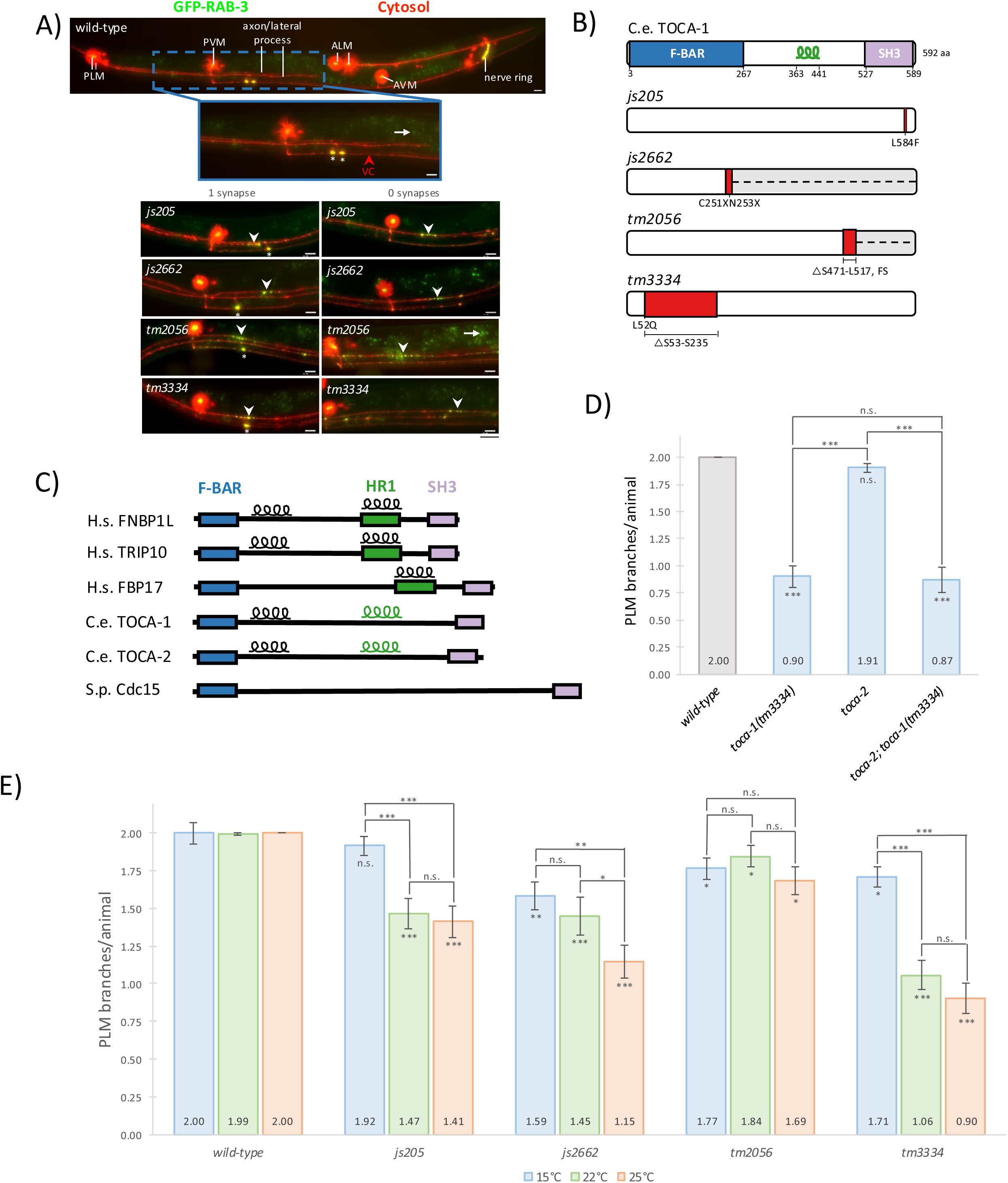
TOCA-1 mutations disrupt PLM synaptic branching in a temperature-sensitive manner. **(A)** Top panel - a representative maximum intensity projections of an L4 larval stage *C. elegans* expressing cytosolic mRFP and synaptic vesicle associated GFP-RAB-3 in TRNs under control of the *mec-7* promoter to visualize the soma and projections of the two PLM neurons, two ALM neurons and the single PVM and AVM neurons. A subpanel focuses on the region in the posterior half of the animal where PLM neurons form a collateral branch to contact the ventral cord (VC) where each branch terminates in a synaptic varicosity. Wild-type animals (top panel) show two distinct PLM branches with large synaptic puncta (white asterisks) in the ventral cord (red arrowhead), while *toca-1* mutants (bottom panel; *js205*, *js2662*, *tm2056*, *tm3334*) display fewer branches and associated synaptic puncta. Additionally, GFP-RAB-3 accumulations are observed in PLM processes lacking branches in the vicinity of the expected branch position (white arrowheads). *C. elegans* accumulate autofluorescent compounds in intestinal gut granules which are visible in some panels. All green signal not co-localizing with red cytosolic signal derives from gut granules (small arrow marks a representative granule). Animals were grown at 25°C and imaged using wide field florescence microscopy. Scale bar=10 μm. **(B)** Schematic representation of *C. elegans* TOCA-1 protein domains and mutant alleles. TOCA-1 contains F-BAR, coiled-coil, and SH3 domains. Mutation sites are indicated: *js205* (C1750T, L584F), *js2662* (C251X, N253X), *tm2056* (ΔS471-L517, frameshift, FS), and *tm3334* (L52Q, ΔS53-S235). Numbers indicate amino acid positions. **(C)** Domain architecture comparison of evolutionarily conserved PCH (<u>p</u>ombe <u>C</u>dc15 <u>h</u>omology) family proteins across species. Human FNBP1L (aka TOCA-1), TRIP10 (aka CIP4), and FNBP1 (aka FBP17), *C. elegans* TOCA-1 and TOCA-2, and *S. pombe* Cdc15 all contain conserved F-BAR and SH3 domains. *C. elegans* TOCA-1 and TOCA-2 have coiled-coil domains with homology to Human TOCA-1 and CIP4 HR1 domains. Adapted from (Ho *et al*., 2004). **(D)** Quantification of PLM branching in *toca-1* and *toca-2* single mutants and *toca-1; toca-2* double mutants. Bar graph shows mean number of PLM branches per animal. *toca-2* mutants show only slight reduction in branching compared to wild-type, and *toca-1(tm3334); toca-2* double mutants are statistically indistinguishable from *toca-1(tm3334)* single mutants. Animals were grown at 25°C and scored as L4 larvae using florescence microscopy. Statistical significance determined by Mann-Whitney U test. n=25-53 animals per condition. Error bars represent SEM. ***p<0.001, n.s.=not significant. **(E)** Temperature sensitivity of PLM branching defects in *toca-1* mutants. Bar graph shows mean number of PLM branches per animal at 15°C, 22°C, and 25°C for wild-type and multiple *toca-1* alleles. Three *toca-1* mutants show increased penetrance of branching defects at higher temperatures, with *tm3334* showing the most severe phenotype at 25°C. Animals were grown at indicated temperatures and scored as L4 larvae using florescence microscopy. Statistical significance determined by Mann-Whitney U test. n=29-81 animals per condition. Error bars represent SEM. ***p<0.001, **p<0.01, *p<0.05, n.s.=not significant.

We isolated *sam-8(js205)* in a clonal screen for animals that failed to form PLM synaptic varicosities (Schaefer, 2001). To characterize the synaptic defect, we first examined PLM morphology using cytosolic markers, which revealed that *sam-8(js205)* mutants displayed reduced or absent collateral branches compared to wild-type animals. We then analyzed synaptic vesicle localization using GFP-RAB-3, finding that wild-type animals show bright puncta in the ventral cord (VC) corresponding to synaptic varicosities, while *sam-8(js205*) mutants exhibited reduced or absent synaptic puncta correlating with the branching defects (Fig. 1A). This synaptic disruption was confirmed using SNB-1::GFP, indicating genuine loss of synaptic structures (Fig. S1A). Analysis with cytosolic PLM markers revealed 100% correlation between loss of PLM branches and absence of VC synaptic puncta.

Notably, PLM processes lacking branches exhibited GFP-RAB-3 accumulation at presumptive branching sites, suggesting that synaptic vesicles are produced but cannot be properly transported due to the absence of branches (Fig. 1A). When branches did form successfully, the resulting synapses were indistinguishable from wild-type based on size and intensity of GFP-RAB-3 signal (Figs. S1B-C), indicating that *sam-8* specifically disrupts branch initiation rather than synaptic maturation. The positions of these branching sites remained constant regardless of whether branches successfully formed, indicating that sam-8 affects the ability to initiate branching rather than the spatial patterning of branch locations (Fig. S1D). Additionally, *sam-8* mutants at the L2 larval stage had the same number of PLM branches as L4 animals (Fig. S1E), further supporting that the defect is specifically in initial branch formation rather than branch maintenance or developmental delay. Importantly, ALM neurons, which form their branches during embryogenesis as part of initial outgrowth rather than as a secondary collateral branching event, showed no branching defects in *sam-8* mutants, confirming the specificity of the phenotype to TRN collateral branch formation (Fig. S1F).

### *sam-8* encodes a Toca-1 homolog

Genetic mapping positioned the causative mutation on the left arm of chromosome X, and whole genome sequencing identified a disruptive L584F (C1750T) lesion in *toca-1,* encoding a member of the pombe Cdc15 homology (PCH) family of proteins. toca-1 is most similar to human FNBP1L/Toca-1 (transducer of Cdc42-dependent actin assembly) which was previously implicated in regulation of actin assembly (Ho *et al*., 2004). Three additional alleles confirmed causation: *toca-1(tm3334[L52Q,ΔS53-S235])*, which removes a significant portion of the N-terminal region, *toca-1(tm2056[ΔS471-L517,FS])* affecting the SH3 domain, and *toca-1(js2662[C251X,N253X])* introducing early stop codons (Fig. 1B). All alleles exhibited similar PLM branching defects (Fig. 1A), confirming that *toca-1* is required for PLM branch formation. We therefore renamed *sam-8* to *toca-1*.

### *toca-2* is not involved in branching

*The C. elegans* genome encodes two FNBP1L/Toca-1 homologs, *toca-1* and *toca-2*. Both proteins share a conserved domain architecture including an F-BAR domain for membrane phospholipid binding, a coiled-coil domain with homology to HR1 domains that binds small GTPases, and an SH3 domain that binds proline-rich sequences in proteins such as WASP (Fig. 1C). To test whether TOCA-2 contributes to PLM branching through functional redundancy, we analyzed *toca-2* single mutants and *toca-1; toca-2* double mutants. *toca-2* mutants showed only slight reduction to 1.91 branches per animal (n.s. compared to wild-type) and *toca-1; toca-2* double mutants were statistically indistinguishable from *toca-1* single mutants (Figs. 1D, S2). This demonstrates that the branching defect is specifically due to TOCA-1 loss rather than combined loss of both paralogs.

### *toca-1* branching defects are temperature sensitive

We initially observed variable penetrance of the *toca-1* branching defect. Further analysis revealed that the *toca-1* PLM branching defect is temperature sensitive, being more penetrant when animals are grown at 25°C, and much less penetrant when animals are grown at 15°C (Fig. 1E). Wild-type animals maintained consistent branch numbers across temperatures, while three *toca-1* alleles showed increased branching defects at higher temperatures, with *tm3334* exhibiting the most severe phenotype at 25°C.

### TOCA-1 localizes to the site of branch formation and functions in the presynaptic cell to regulate branching

Tissue-specific rescue experiments distinguished between cell-autonomous and non-cell-autonomous functions. Expression under the *mec-4* promoter (active in PLM) fully rescued the branching defect in *toca-1(tm3334)* mutants, while expression under the *glr-1* promoter (active in interneurons postsynaptic to PLM, but not in PLM) failed to rescue (Figs. 2A-B). These results demonstrate that TOCA-1 functions cell-autonomously within presynaptic PLM neurons.

**Figure 2.**
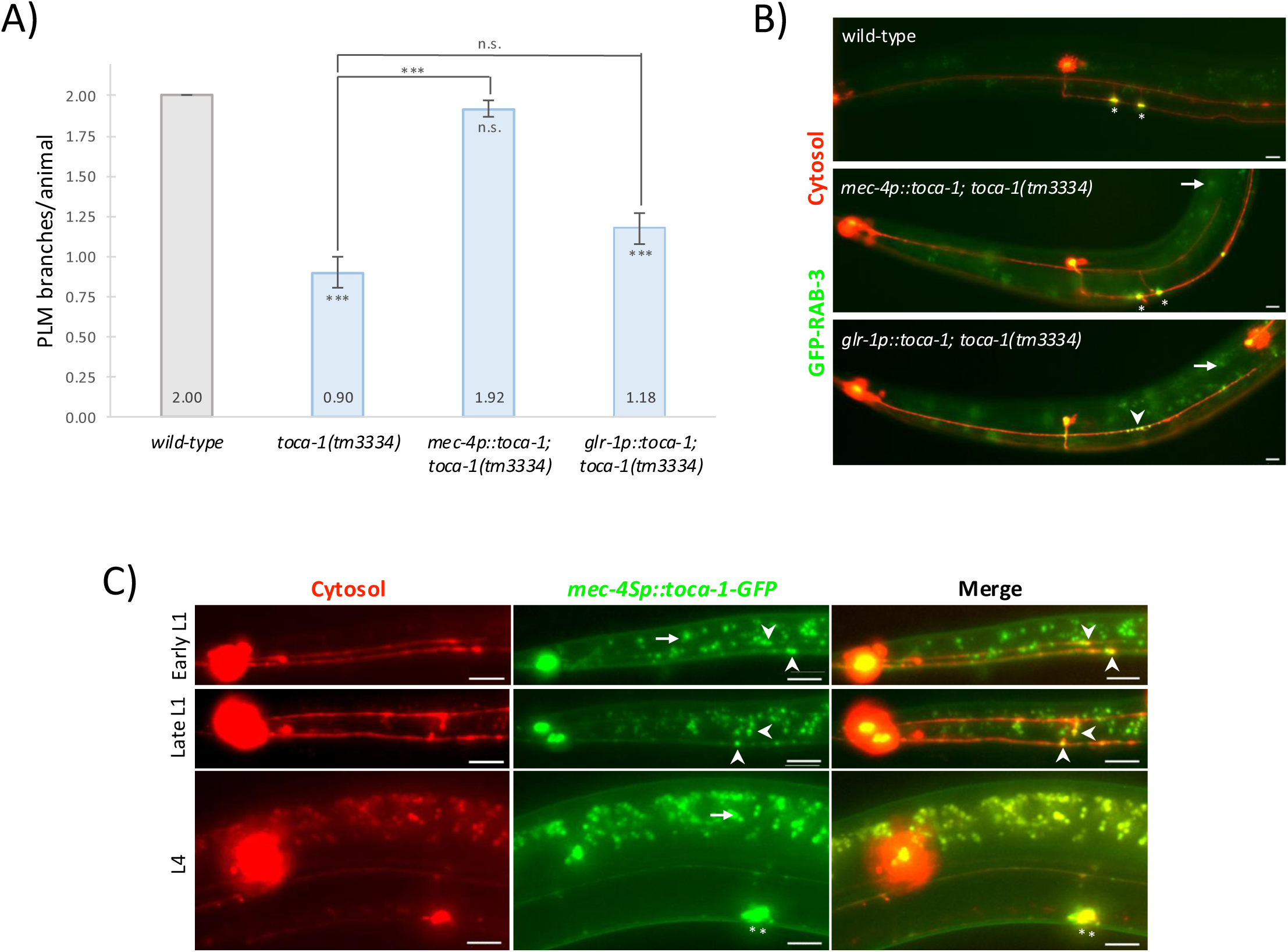
TOCA-1 functions cell-autonomously in PLM neurons and localizes to branch formation sites. **(A)** Quantification of rescue experiments demonstrating cell-autonomous function of TOCA-1. Bar graph shows mean number of PLM branches per animal. Animals expressing *toca-1* under the TRN-specific *mec-4* promoter restore normal branching levels in *toca-1(tm3334)* mutants, while animals expressing *toca-1* in interneurons post-synaptic to PLM under the *glr-1* promoter shows no rescue effect. Animals were grown at 25°C and scored as L4 larvae using florescence microscopy. Statistical significance determined by Mann-Whitney U test. n=25-57 animals per condition. Error bars represent SEM. ***p<0.001, n.s.=not significant. **(B)** Representative images showing tissue-specific rescue of *toca-1(tm3334)* branching defects. Expression of TOCA-1 under the *mec-4* promoter fully rescues branching as indicated by normal branch number and synaptic puncta (asterisks), while expression under the *glr-1* promoter fails to rescue branching defects and retains the GFP-RAB-3 accumulation along the PLM process (arrowhead). Images are maximum intensity projections. Scale bar=10 μm. **(C)** Subcellular localization of TOCA-1::GFP in PLM neurons in maximum intensity projections. TOCA-1::GFP forms puncta along the PLM process with enrichment at branch formation sites (arrowheads) and in synaptic puncta (asterisks), consistent with a direct role in branch initiation. Signal in the green channel that does not co-localize with the PLM-derived red cytosolic signal is autofluorescence from gut granules. Animals were grown at 22°C and imaged using florescence microscopy. Scale bar=10 μm.

To determine the subcellular localization of TOCA-1 in PLM neurons, we expressed a TOCA-1::GFP fusion protein under the control of the *mec-4* promoter in a strain with a cytosolic marker. TOCA-1::GFP expressed under *mec-4* promoter formed puncta along the PLM process with enrichment at branch formation sites (Fig. 2C), consistent with TOCA-1 playing a direct role in initiating branch formation. In the wild type, PLM branches initiate about 2 hours after hatching. In early L1s we observed accumulation of TOCA-1::GFP in the vicinity where branch formation initiates about 2/3 of the length down the process. As branch formation initiates TOCA-1::GFP is located very close to the branch protrusion and continues to localize near the branch tip as the branch extends toward the ventral cord. In older animals, TOCA-1::GFP remains concentrated at the synaptic varicosity but is much less abundant along the length of the process. Thus, TOCA-1 is properly positioned to play an instructive role in branch initiation and extension.

### *toca-1* functions in conjunction with *cdc-42* to regulate branching

Cdc42 plays a critical role in regulating actin dynamics functioning in conjunction with Wiskott-Aldrich syndrome protein (WASP) and TOCA family proteins (Watson *et al*., 2017). TOCA-1 contains an HR1 domain that binds Cdc42, integrating Cdc42 signaling with actin cytoskeleton dynamics (Ho *et al*., 2004; Lee *et al*., 2010). Similarly, *C. elegans toca-1* has been documented to integrate worm *cdc-42* signaling in multiple contexts (Giuliani *et al*., 2009; Ouellette *et al*., 2016; Raduwan *et al*., 2020). Examination of *C. elegans cdc-42(gk388)* null mutants revealed significant reduction in PLM branching, similar to *toca-1* mutants (Figs. 3A-B). Furthermore, in wild-type animals, TRN-specific expression of dominant-negative *cdc-42(T17Ndn)* caused severe branching defects, gain-of-function *cdc-42(G12Vgf)* caused even more severe reduction, while over-expression of wild-type CDC-42 had no effect. This indicates that both loss and hyperactivation of CDC-42 disrupt proper branching.

**Figure 3.**
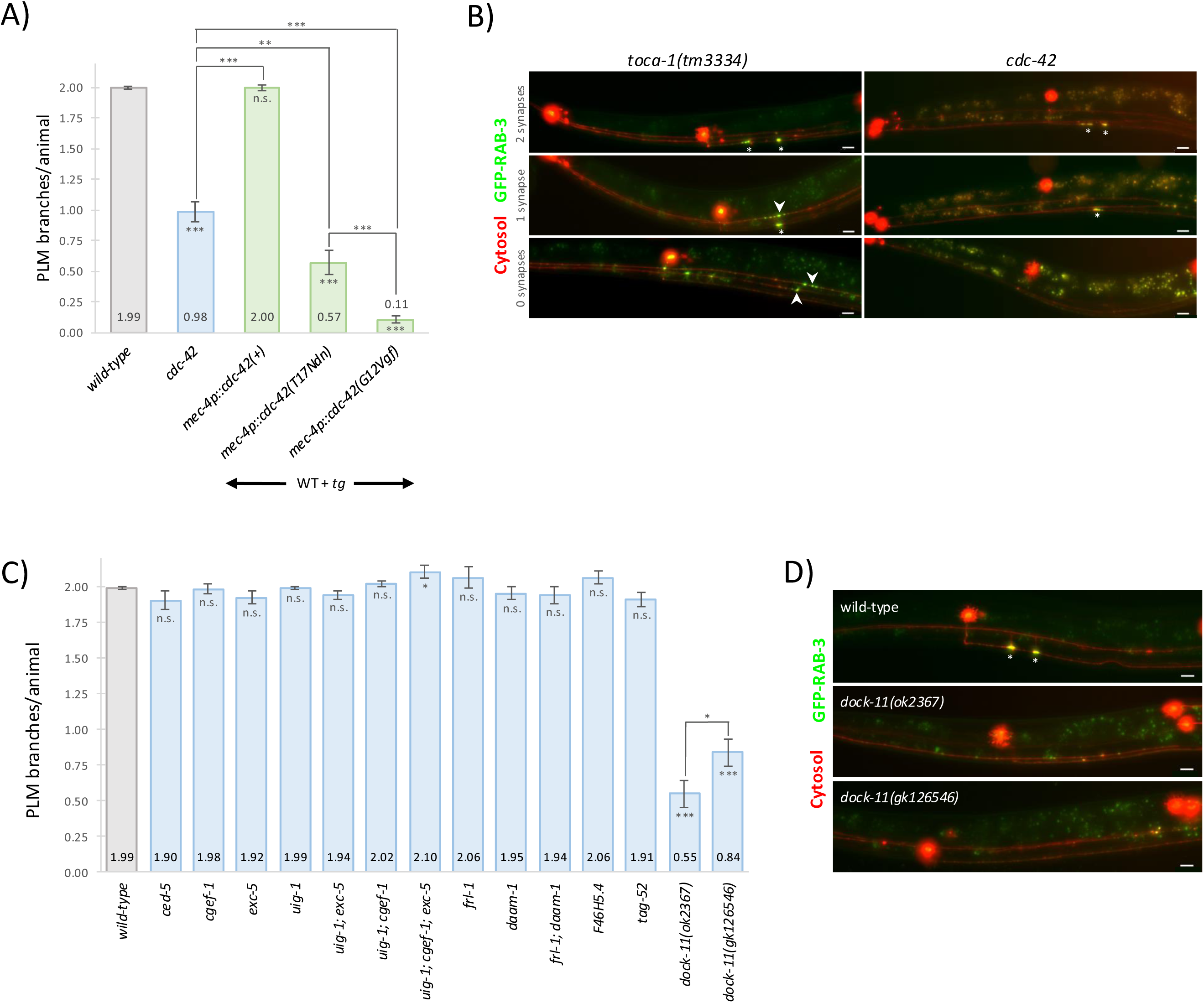
CDC-42 and DOCK-11 GEF regulate PLM branching in conjunction with TOCA-1. **(A)** Quantification of PLM branching in various *cdc-42* alleles. Bar graph shows mean number of PLM branches per animal. Animals were grown at 22°C and scored as L4 larvae using fluorescence microscopy. *cdc-42(gk388)* mutants show significant reduction in PLM branching. TRN-specific expression of dominant-negative *cdc-42(T17Ndn)* and gain-of-function *cdc-42(G12Vgf)* both cause severe branching defects when expressed in genomic *cdc-42(+)* background, but overexpression of wild-type *cdc-42* had no effect. This indicates that proper *cdc-42* regulation is critical for normal branching. Statistical significance determined by Mann-Whitney U test. n=42-118 animals per condition. Error bars represent SEM. ***p<0.001, **p<0.01, n.s.=not significant. **(B)** Maximum intensity projection fluorescent images of PLM neurons in *toca-1* and *cdc-42(gk388)* mutants showing representative animals with 2, 1, and 0 synapses (asterisks). Arrows indicate accumulation of GFP-RAB-3 at presumptive branching sites in *toca-1* animals when branches fail to form. This characteristic GFP-RAB-3 accumulation phenotype is not observed in *cdc-42* animals. Scale bar=10 μm. **(C)** Screen of candidate CDC-42 GEFs for PLM branching defects. Bar graph shows mean number of PLM branches per animal. Animals were grown at 22°C and scored as L4 larvae. Most GEF mutants show normal PLM branching, while two independent derived *dock-11* mutants showed the severe reduction in branching. Statistical significance determined by Mann-Whitney U test. n=21-104 animals per condition. Error bars represent SEM. ***p<0.001, *p<0.05, n.s.=not significant. **(D)** Representative maximum intensity projection fluorescent images of PLM neurons in *dock-11* mutants showing branching defects similar to those of *cdc-42* mutants, confirming therequirement for DOCK-11 in PLM branch formation. As is the case with *cdc-42* mutants, *dock-11* do not show the characteristic GFP-RAB-3 accumulations in the vicinity of the expected branch site when branches are absent. Scale bar=10 μm.

To identify the guanine nucleotide exchange factor (GEF) responsible for CDC-42 activation during branching, we examined mutants disrupting several candidate *C. elegans* Rho family GEFs (Chan and Nance, 2013). Most GEF mutants including *ced-5*, *cgef-1*, *exc-5*, and *uig-1* showed normal PLM branching or mild defects (Fig. 3C). However, *dock-11(ok2367)* showed significantly reduced branching, confirmed by a second independent allele *dock-11(gk126546)* (Figs. 3C-D). This identifies DOCK-11 as a critical component of the PLM branching machinery likely acting as a CDC-42 GEF. Additionally, analysis of GFP-RAB-3 localization revealed distinct patterns among the mutants. In unbranched PLM processes of *toca-1* animals RAB-3::GFP accumulations were consistently observed near the expected branching site, whereas *cdc-42* and *dock-11* mutants showed dramatically reduced RAB-3 accumulations (Fig. S3). This suggests that CDC-42 and DOCK-11 function upstream to specify branching sites and also modulate vesicle trafficking, while TOCA-1 acts downstream to execute branch formation after vesicles have been recruited to presumptive branching sites.

### *toca-1* functions in conjunction with *wsp-1* to regulate branching

In vertebrate systems, TOCA-1 functions as a scaffold protein that binds both Cdc42 and N-WASP, facilitating their interaction and enhancing actin nucleation (Ho *et al*., 2004). WSP-1 is the *C. elegans* ortholog of human WASP, containing seven conserved functional domains including the WH1 domain and the C-terminal VCA (Verprolin Central Acid) domain that is essential for activating the Arp2/3 complex and promoting actin polymerization (Kurisu and Takenawa, 2009) (Fig. 4A). The V region binds actin monomers and the CA region binds the Arp2/3 complex; together they stimulate rapid actin polymerization. In the absence of Cdc42^GTP^, WASP is auto-inhibited. In *C. elegans*, TOCA-1 has been shown to bind both WSP-1 and CDC-42^GTP^ for proper actin organization and membrane trafficking (Giuliani *et al*., 2009).

**Figure 4.**
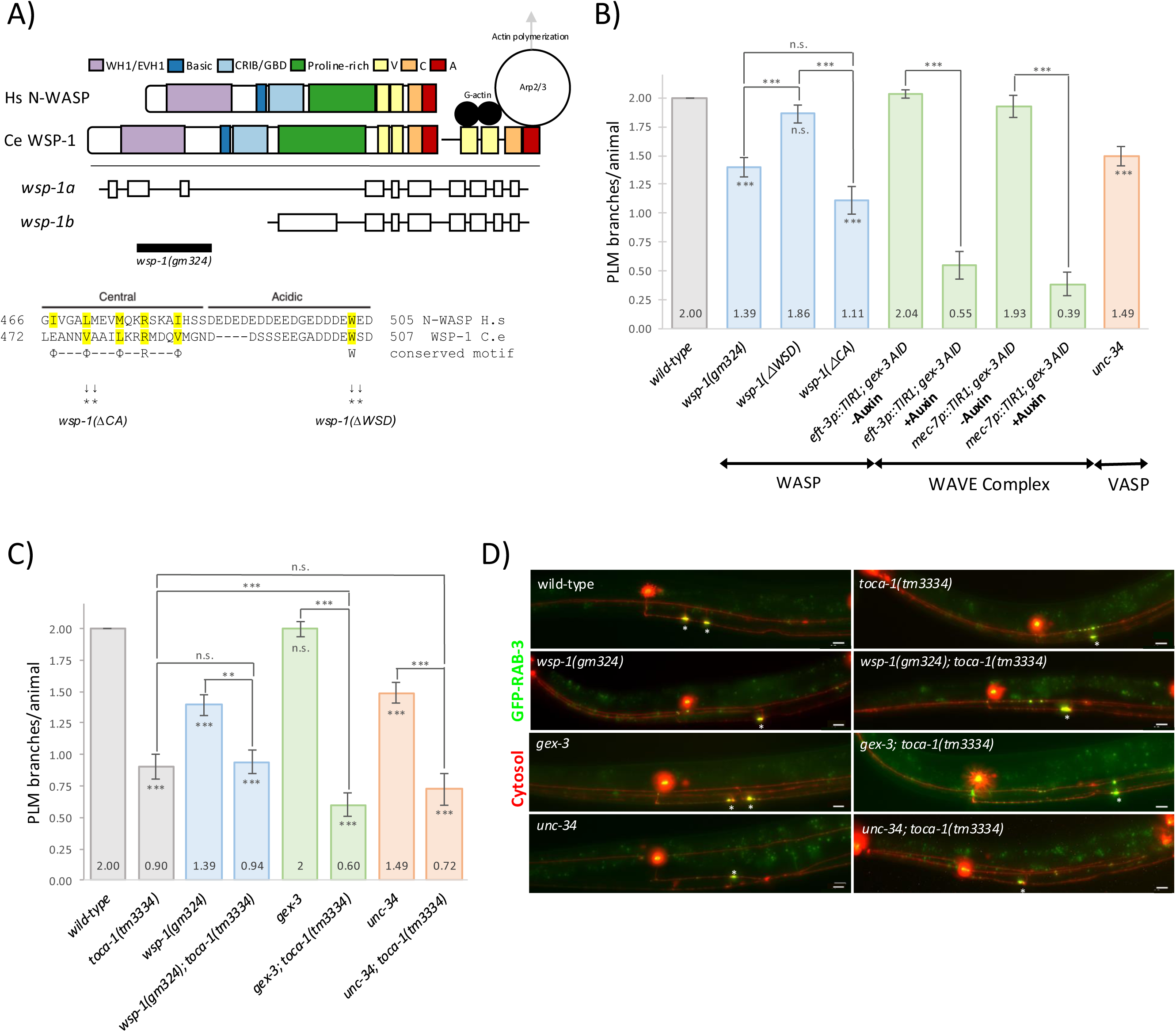
WSP-1 and WAVE complex components regulate PLM branching. **(A)** Schematic of WSP-1 protein domains and mutant alleles. WSP-1 contains multiple conserved functional domains with Hs. N-WASP including the C-terminal VCA region. Two alleles target specific regions: ΔWSD, affecting only the acidic domain, and ΔCA, affecting both the central and acidic domains. The conserved residues in are highlighted in yellow. R=Arg, W=Trp, X=any residue, `Φ`=long chain aliphatic residue. The *gm324* allele only targets one the *wsp-1a* isoform. Adapted from (Panchal *et al*., 2003) and (Kurisu and Takenawa, 2009). **(B)** Quantification of PLM branching in actin regulatory mutants. Bar graph shows mean number of PLM branches per animal. Animals were grown at 25°C and scored as L4 larvae using florescence microscopy. Analysis of WASP function reveals that *wsp-1* shows substantially reduced branching. Domain-specific deletions demonstrate that ΔCA causes the most severe defect, more severe than both *wsp-1* and ΔWSD, indicating the critical importance of the complete CA region for robust WASP function in branching. WAVE complex involvement is demonstrated by severe branching reduction following GEX-3 AID depletion. *unc-34* mutants show moderate branching reduction, indicating the importance of VASP. Statistical significance determined by Mann-Whitney U test. n=25-66 animals per condition. Error bars represent SEM. ***p<0.001, n.s.=not significant. **(C)** Double mutant analysis reveals genetic relationships. Animals were grown at 25°C and scored as L4 larvae. *wsp-1; toca-1* and *unc-34; toca-1* double mutants show branching defects indistinguishable from *toca-1* single mutants. By contrast, toca-1; *gex-3(av259[L353F])* hypomorph double mutants show a reduction compared to either single mutant suggesting the WAVE complex acts in parallel to toca-1. Statistical significance determined by Mann-Whitney U test. n=25-66 animals per condition. Error bars represent SEM. ***p<0.001, **p<0.01, n.s.=not significant. **(D)** Representative maximum intensity projections fluorescent images of PLM neurons in single mutants in the actin regulatory proteins [*wsp-1(gm324)*, *unc-34(e315)*, *gex-3(av259)*] and *toca-1(tm3334)* double mutants probing the genetic relationships between actin regulatory pathways and TOCA-1 in PLM branching. Asterisks indicate synaptic puncta. Scale bar=10 μm.

The *wsp-1(gm324)* allele, which specifically affects the *wsp-1a* isoform, exhibited substantially reduced branching (Figs. 4B, S4A). To determine whether the Arp2/3 recruitment activity of WSP-1 was required for branch formation, we generated two additional alleles: *wsp-1(js1377)* with deletion in the A region (ΔWSD) and *wsp-1(js1380)* with a deletion truncating the CA domain (ΔCA) (Fig. 4A). The ΔCA deletion caused a severe branching defect, indicating that the complete CA region is essential for promoting branching (Figs. 4B, S4A).

### *arp-2* is required for branch formation, but not extension of non-branching processes

Given the role of WSP-1 in activating the Arp2/3 complex, we investigated the requirement of Arp2/3 components in PLM branching. We examined *arx-5*, which encodes the ARP3 subunit, and *arx-2*, which encodes the ARP2 subunit of the Arp2/3 complex. Both mutants are homozygous lethal with animals arresting as sterile adults, and progeny homozygous for these mutations exhibited normal PLM branching, likely due to maternal rescue (Figs. S5A-B). RNAi targeting of *arx-5* also failed to produce branching defects (Fig. S5A-B). To eliminate the maternal contribution, we tagged the endogenous *arx-2* gene with an auxin-inducible degron tag (Zhang *et al*., 2015). When combined with a ubiquitously expressed TIR1 auxin response gene, addition of auxin led to embryonic or early larval arrest depending on whether auxin was added before or after fertilization. Branching was disrupted in the arrested larvae (Figs 5A-B). When TIR1 was restricted to touch receptor neurons using a *mec-7* promoter, *arx-2 AID* strains showed severe branching reduction upon auxin treatment, with complete loss of branches in TRN-specific TIR1 animals (Figs. 5A-B). This demonstrates that ARX-2 is essential for PLM branch formation.

**Figure 5.**
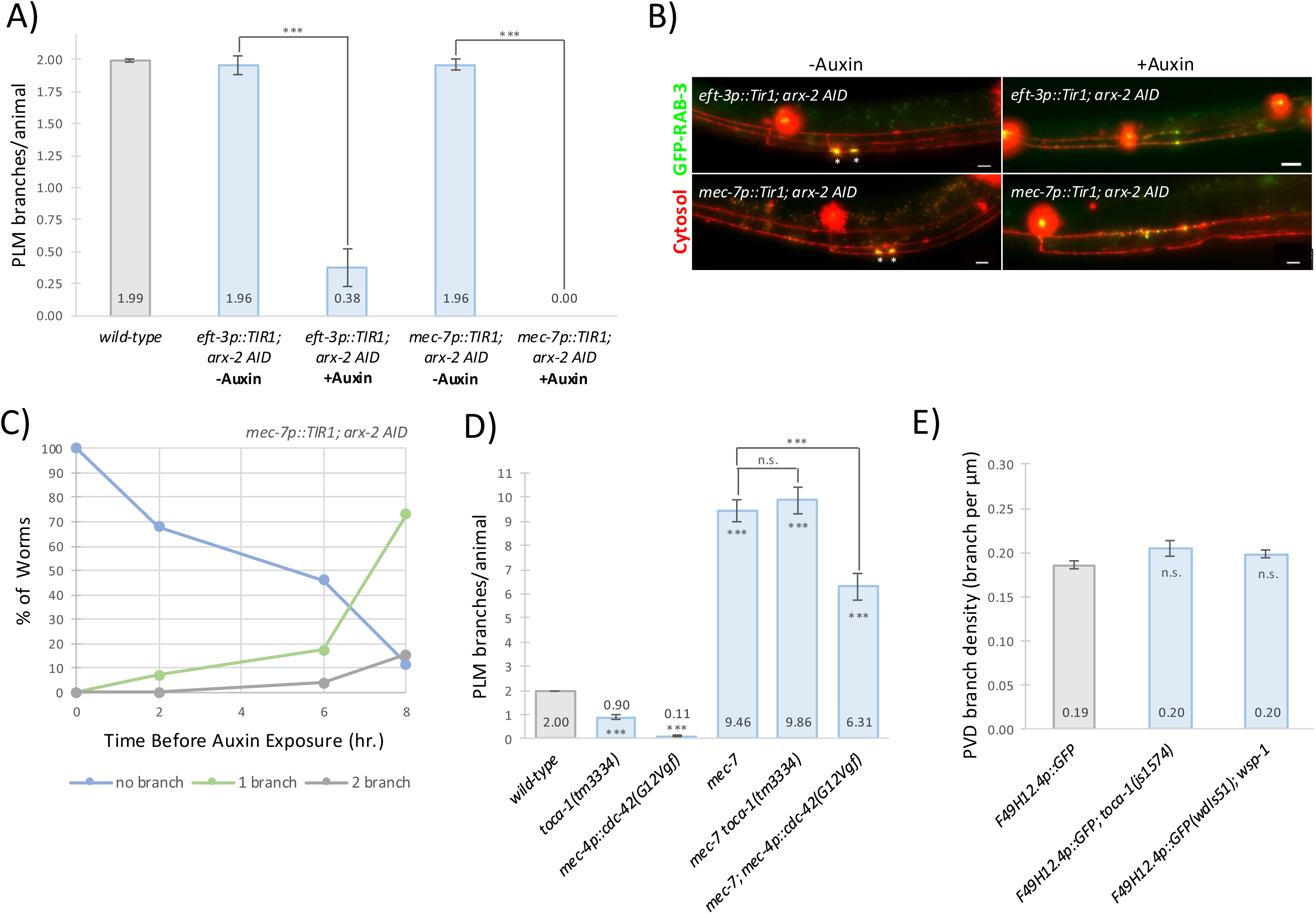
ARX-2/ARP2 is required for PLM branch initiation but not maintenance. **(A)** Quantification of PLM branching in *arx-2* AID strains. Bar graph shows mean number of PLM branches per animal. Animals were grown at 22°C and scored as L4 larvae. Auxin treatment results in severe reduction in branching, with complete loss of branches in TRN-specific TIR1 animals, demonstrating that the Arp2/3 complex is essential for PLM branch formation. Statistical significance determined by Mann-Whitney U test. n=24-50 animals per condition. Error bars represent SEM. ***p<0.001. **(B)** Representative maximum intensity projection fluorescent images of PLM neurons in *arx-2* AID strains with and without auxin treatment, showing dramatic loss of branches upon ARX-2 depletion. I Scale bar=10 μm. **(C)** Temporal analysis of ARX-2 requirement in PLM branching shows the percentage of animals with PLM branches when auxin treatment is initiated at different developmental time points (immediately after hatching vs. later exposure). This demonstrates that ARX-2 is required during branch initiation but not for branch maintenance. Animals were grown at 22°C with auxin treatment starting at indicated time points and scored as L4 larvae. By 8 hours post-hatching, approximately 70% developed one branch and 15% developed two branches. n=26-57 animals per condition. **(D)** Analysis of TOCA-1 function in aberrant hyper-branching. *mec-7* mutants exhibit disorganized hyper-branching that is unaffected by a *toca-1* mutation, while *cdc-42(G12Vgf)* partially suppresses the hyper-branching phenotype. Animals were grown at 25°C and scored as L4 larvae. Statistical significance determined by Welch’s t-test. n=25-113 animals per condition. Error bars represent SEM. ***p<0.001, n.s.=not significant. **(E)** Quantification of mean PVD branch density (branches per unit length of dendrite) for wild-type, *toca-1*, and *wsp-1* mutants. The results demonstrate that there is no difference in PVD neuron density, visualized with PVD cell-specific GFP reporter (F49H12.4::GFP). Animals were grown at 25°C and scored as L4 larvae. Statistical significance determined by Welch’s t-test. n=42-60 animals per condition. Error bars represent SEM. n.s.=not significant.

Time-course experiments revealed that ARX-2 is required during branch initiation but not maintenance. *mec-7p::TIR1; arx-2 AID* animals exposed to auxin immediately after hatching showed complete branch loss, while later exposure allowed increasing percentages to develop branches. By 8 hours post-hatching, approximately 70% developed one branch and 15% developed two branches (Fig. 5C).

### *toca-1* is specifically required for stereotyped collateral branching in PLM neurons

We tested whether TOCA-1 functions as a general branching regulator or plays a more specific role in PLM collateral branch formation. To address this question, we examined two distinct branching contexts, aberrant hyper-branching and normal dendritic arbor formation.

The *mec-7* gene encodes a β-tubulin essential for mechanosensory neuron development and *mec-7* mutants exhibit disorganized hyper-branching that lacks the stereotyped positioning of normal PLM collateral branches (Figs. 5D, S6A). This hyper-branching was unaffected in *mec-7 toca-1* double mutants, demonstrating that excessive branching occurs through a TOCA-1-independent mechanism. In contrast, *mec-7; cdc-42(G12Vgf)* double mutants showed partial suppression, indicating CDC-42 can modulate hyper-branching through pathways distinct from TOCA-1.

Second, we examined PVD sensory neurons, which form elaborate dendritic arbors. Neither *toca-1(js1574)* nor *wsp-1* affected PVD branch density (Figs. 5E, S6B), despite causing dramatic PLM branching defects. These results suggest that TOCA-1 plays a specific role in PLM collateral branch formation rather than serving as a universal branching regulator.

### Role of WAVE complex components and UNC-34/Ena/VASP in PLM branching

Since Arp2/3 is 100% essential for branching while *toca-1*, *cdc-42*, and *wsp-1* mutants only show partial defects, we examined whether other pathways known to activate the Arp2/3 complex may also play a role in PLM branch formation. We investigated the WAVE complex, another major activator of Arp2/3 (Kurisu and Takenawa, 2009). We examined several WAVE complex components including *wve-1* (encoding the WAVE protein), *gex-2*, and *gex-3*.

*wve-1* and *gex-2* homozygous progeny from balanced heterozygotes showed normal PLM branching, likely due to maternal rescue. *gex-3* mutants also showed normal branching (Figs. S4A-B). However, auxin-inducible degron (AID) targeting of GEX-3 in animals with ubiquitous or TRN-specific TIR1 expression showed severe branching reduction (Figs. 4B, S4A), demonstrating WAVE complex involvement in PLM branching.

We also examined *unc-34*, which encodes the *C. elegans* ortholog of Ena/VASP proteins that regulate actin dynamics independently of the Arp2/3 complex (Withee et al., 2004). *unc-34* mutants exhibited moderate reduction in PLM branching (Figs. 4B, S4A), indicating that PLM branching requires both Arp2/3-mediated actin nucleation and UNC-34-mediated actin regulation.

Double mutant analysis revealed distinct genetic relationships. *wsp-1; toca-1* and *unc-34; toca-1* double mutants showed branching defects indistinguishable from *toca-1* single mutants, indicating complete epistasis (Figs. 4C-D). *gex-3; toca-1* double mutants exhibited a reduction in branch formation compared to either single mutant, showing that the WAVE complex provides some independent contribution alongside the TOCA-1/WSP-1 pathway.

### TOCA-1 affects branching without disrupting mitochondrial or synaptic protein localization

To determine whether TOCA-1 has broader effects on neuronal organization beyond branching, we examined the distribution of mitochondria and active zone components along the PLM process in *toca-1* mutants. Using mitochondrial (*mec-7p::mito-GFP*) and active zone protein (*mec-7p::GFP-ELKS-1*) markers, we found that mitochondrial density, GFP-ELKS-1 density, and GFP-ELKS-1 fluorescence were comparable between wild-type and *toca-1* mutants (Figs. 7SA-C). This indicates that TOCA-1 functions specifically in branch initiation rather than general cellular organization.

### Role of netrin signaling in PLM branching

To investigate potential guidance cues involved in PLM branch formation, we examined components of the netrin signaling pathway, which is known to regulate axon guidance and branching (Kennedy *et al*., 1994; Serafini *et al*., 1994; Dent *et al*., 2004). Single mutants *unc-5*, *unc-6*, and *unc-40* each showed modest reductions in PLM branching (Figs. 6A-B). Double mutants combining *unc-5, unc-6* or *unc-40* with *toca-1(tm3334)* showed a slight further reduction in branching compared to *toca-1* single mutants (Figs. 6A-B), indicating that netrin signaling contributes to PLM branch formation. Both PVM and AVM neurons showed increased morphological abnormalities in netrin pathway mutants, with double mutants combining netrin components with *toca-1* showing enhanced defects compared to netrin single mutants (Figs. 6C-D, S8). This enhancement suggests that TOCA-1 and netrin pathways converge on shared downstream effectors that regulate neuronal morphology, despite TOCA-1’s primary specificity for PLM collateral branching.

**Figure 6.**
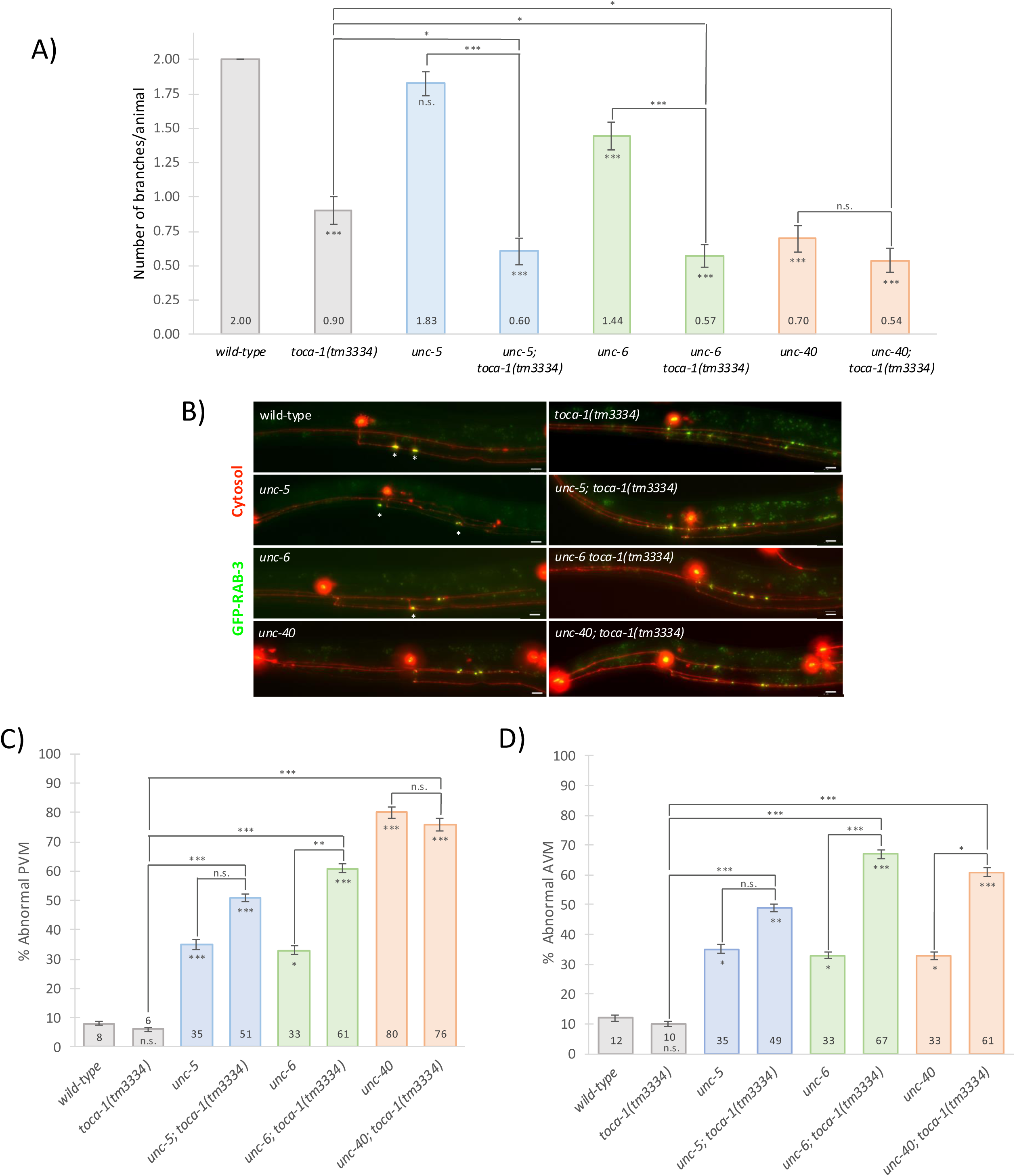
Netrin signaling contributes to PLM branch formation in the same pathway as TOCA-1. **(A)** Quantification of PLM branching defects in netrin pathway mutants. Bar graph shows mean number of PLM branches per animal for wild-type, single mutants, *toca-1(tm3334)* single mutant, and double mutants combining netrin pathway genes with *toca-1*. Animals were grown at 25°C and scored as L4 larvae using florescence microscopy. Single mutants *unc-5*, *unc-6*, and *unc-40* each show reductions in PLM branching compared to wild-type. Double mutants combining *unc-5, unc-6* or *unc-40* with *toca-1* show a reduction in branching compared to *toca-1* single mutants. Statistical significance determined by Mann-Whitney U test. n=39-61 animals per condition. Error bars represent SEM. ***p<0.001, *p<0.05, n.s.=not significant. **(B)** Representative images of PLM neurons expressing GFP-RAB-3 and cytosolic mRFP in netrin signaling mutants. Asterisks indicate synaptic puncta. Images are maximum intensity projections. Scale bar=10 μm. **(C)** Quantification of PVM morphological abnormalities in netrin pathway mutants. Bar graph shows percentage of animals with abnormal PVM morphology for wild-type, single mutants, *toca-1(tm3334)* single mutant, and double mutants combining netrin pathway genes with *toca-1.* Animals were grown at 25°C and scored as L4 larvae using fluorescence microscopy. Abnormal phenotypes include failure to meet the ventral nerve cord, absence of the neuron, incorrect process length, branching defects, migration paths with >45 degree turns, or ventral growth patterns (categories shown in Figure S8). Single mutants *unc-5*, *unc-6*, and *unc-40* each show increased abnormal morphology compared to wild-type. The *unc-6; toca-1* double mutant shows a statistically significant increase in abnormal phenotypes compared to the respective single mutants. Statistical significance determined by chi-squared test with Yates continuity correction. n=25-61 animals per condition. Error bars represent SD. ***p<0.001, **p<0.01, *p<0.05, n.s.=not significant. **(D)** Quantification of AVM morphological abnormalities in netrin pathway mutants. Bar graph shows percentage of animals with abnormal AVM morphology for wild-type, single mutants, *toca-1(tm3334)* single mutant, and double mutants combining netrin pathway genes with *toca-1.* Animals were grown at 25°C and scored as L4 larvae using fluorescence microscopy. Abnormal phenotypes scored as described for PVM neurons (Figure S8). Single mutants *unc-5*, *unc-6*, and *unc-40* each show increased abnormal morphology compared to wild-type. The *unc-6; toca-1* and *unc-40; toca-1* double mutants show statistically significant increases in abnormal phenotypes compared to the respective single mutants. Statistical significance determined by chi-squared test with Yates continuity correction. n=25-61 animals per condition. Error bars represent SD. ***p<0.001, **p<0.01, *p<0.05, n.s.=not significant.

## Discussion

We have identified TOCA-1 as an essential organizer of PLM collateral branch formation, revealing molecular components that coordinate membrane dynamics with actin nucleation to achieve precise neuronal connectivity. Our findings establish that TOCA-1 functions cell-autonomously within PLM neurons and requires DOCK-11, CDC-42, WSP-1, and the Arp2/3 complex for proper function, providing mechanistic insights into one of the most stereotyped examples of neuronal branching in *C. elegans*.

### TOCA-1 as a spatial organizer of neuronal morphogenesis

Our work reveals a novel role for TOCA-1 as an essential organizer required for neuronal branching. While previous studies established TOCA-1’s biochemical interactions with CDC-42 and WASP proteins (Ho *et al*., 2004; Bu *et al*., 2009), our findings demonstrate that TOCA-1 function is required for normal branching alongside multiple actin regulatory pathways (WSP-1, WAVE complex, and UNC-34). The enrichment of TOCA-1 at branch initiation sites, combined with the requirement for DOCK-11-mediated CDC-42 activation, reveals that spatial control may emerge from precisely localized assembly of multi-component branching machinery.

The temperature sensitivity of branching defects provides insight into the precision required for this process, similar to other genes affecting synaptic development (Gu *et al*., 1996; Schaefer *et al*., 2000). Like other developmentally critical processes that become disrupted when cellular kinetics are altered, PLM branch formation appears to require temporal coordination that becomes compromised at elevated temperatures. This suggests that the window for successful branch initiation is narrow and depends on precise timing of molecular interactions.

### TOCA-1 function in filopodia formation across biological systems

Our findings place PLM branching within the broader context of both general axon branching principles and TOCA-1’s specific molecular functions across biological systems. Extensive studies by Gallo and colleagues have established that axon collateral branches are initiated by actin filament-based axonal protrusions that subsequently become invaded by microtubules, with filopodia emerging from transient actin patches that serve as cytoskeletal precursors (Gallo, 2011; Gallo, 2013). These axonal actin patches require the Arp2/3 complex for their formation and contain regulatory proteins such as WAVE (Ketschek and Gallo, 2010), 2010; (Spillane *et al*., 2011).

TOCA-1 functions as a molecular organizer that coordinates the machinery underlying these general principles. TOCA-1 directly binds both CDC42 and N-WASP, serving as a bridge that couples small GTPase signaling to Arp2/3-mediated actin polymerization (Ho *et al*., 2004; Bu *et al*., 2009). TOCA-1 induces filopodia formation through its F-BAR domain for membrane binding, HR1 domain for CDC42 interaction, and SH3 domain for N-WASP recruitment (Bu *et al*., 2009). Toca-1 is also capable of inducing filapodial-like structures in *in vitro* actin assembly assays (Lee *et al*., 2010). Recent analysis has revealed that TOCA-1 clusters promote filopodial protrusion through density-dependent interactions with Ena/VASP proteins (Blake *et al*., 2024).

Our demonstration that TOCA-1 enriches at PLM branch initiation sites and requires DOCK-11, CDC-42, WSP-1, and Arp2/3 for function connects these findings to PLM neurons. PLM branching employs the conserved actin patch-to-filopodia mechanism (Chen *et al*., 2017) while achieving the spatial precision required for stereotyped connectivity through TOCA-1’s role as a molecular organizer that coordinates membrane dynamics with cytoskeletal remodeling.

### Distinct molecular mechanisms underlie different types of neuronal branching

The specificity of TOCA-1’s requirement for PLM collateral branching, without effects on PVD dendritic branching or aberrant hyper-branching, demonstrates that neurons employ distinct molecular toolkits for different morphogenetic programs. This finding challenges the assumption that core cytoskeletal components function universally across branching contexts. Instead, our results suggest that while fundamental processes like actin nucleation may be shared, their regulatory mechanisms are highly specialized.

The contrast with PVD dendritic branching is particularly striking given that both processes involve Arp2/3-mediated actin nucleation (Zou *et al*., 2018), yet only PLM branching requires TOCA-1. This implies that different F-BAR domain proteins or alternative membrane-actin coordinators may operate in distinct cellular contexts, potentially explaining how neurons can simultaneously employ multiple branching programs without interference.

### Integration of intrinsic and extrinsic regulatory mechanisms

Our findings suggests that PLM branch formation requires integration of multiple regulatory inputs. The relationship between netrin signaling components and TOCA-1 demonstrates that guidance cues do not function as independent pathways but instead may provide spatial information that may influence branch formation, consistent with studies showing that axon guidance molecules also affect branching decisions (Yates *et al*., 2001; Dent *et al*., 2004). The Wnt-planar polarity pathway also has previously been documented to influence the AP position of PLM branches (Chen *et al*., 2017). By contrast, loss of TOCA-1 has no influence on the AP position of branch formation suggesting TOCA-1 may act downstream of integration of spatial information.

### Implications for concentration-dependent assembly mechanisms

Our findings suggest that TOCA-1-mediated PLM branching may operate through threshold-based activation mechanisms, providing a solution to a fundamental challenge in neural development: how to achieve precise spatial patterning without relying on complex gene expression programs or elaborate guidance molecule gradients. The enrichment of TOCA-1 at specific branch initiation sites, combined with its requirement for multiple interacting partners including DOCK-11, CDC-42, WSP-1, and Arp2/3, suggests that neurons could utilize intrinsic concentration-dependent switches that activate only when sufficient molecular components accumulate at appropriate locations. Such concentration-dependent thresholds may represent a general strategy for achieving spatial precision in developmental processes, similar to morphogen gradients that control cell fate decisions. The identification of this mechanism in the simple PLM branching system provides an accessible model for understanding how spatial information is encoded in molecular interactions.

### Recruitment of SVs at branch site

Multiple cellular events are required for forming a collateral branch. Filopodia must be induced at the branching site, but in additional, synaptic vesicular components also are concentrated near the branch site. It remains uncertain whether these vesicles participate in the delivery of the membrane components required from filopodial extension, or if these vesicles are being sequestered to facilitate assembly of the nascent synapse developing at the end of the branch once it enters the ventral nerve cord. Regardless of their role, differences in phenotypes in the mutants we analyzed suggest that *dock-11 and cdc-42* mutants disrupt both branch formation and vesicle recruitment while *toca-1* and all the other mutants we analyzed disrupt only the downstream filapodial assembly process while leaving vesicle recruitment intact. Specifically, *toca-1* mutants exhibit large accumulations rich in SV components in the vicinity of the branch site, while *cdc-42* and *dock-11* mutants do not. This is consistent with CDC-42 activation performing two distinct roles 1) recruiting or trapping SVs (or precursors) near the branch site, and 2) promoting filapodial formation. Further work will be required to define the molecular pathways which foster SV accumulation near the branch formation position.

### Model for TOCA-1-mediated branch formation

Based on our findings, we propose a model where TOCA-1 functions as an essential component integrating membrane dynamics with actin regulatory pathways to control PLM collateral branching (Fig. 7). In this model, DOCK-11 acts upstream as a CDC-42 GEF, providing spatial regulation that determines where CDC-42 becomes activated along the PLM axon. TOCA-1 then serves as a crucial organizing protein that is required for multiple downstream effectors: its F-BAR domain induces membrane curvature at branch initiation sites while its coiled-coil and SH3 domains recruit and organize CDC-42 and WSP-1 to drive localized Arp2/3-mediated actin nucleation, consistent with biochemical studies in other systems (Ho *et al*., 2004; Bu *et al*., 2009).

**Figure 7.**
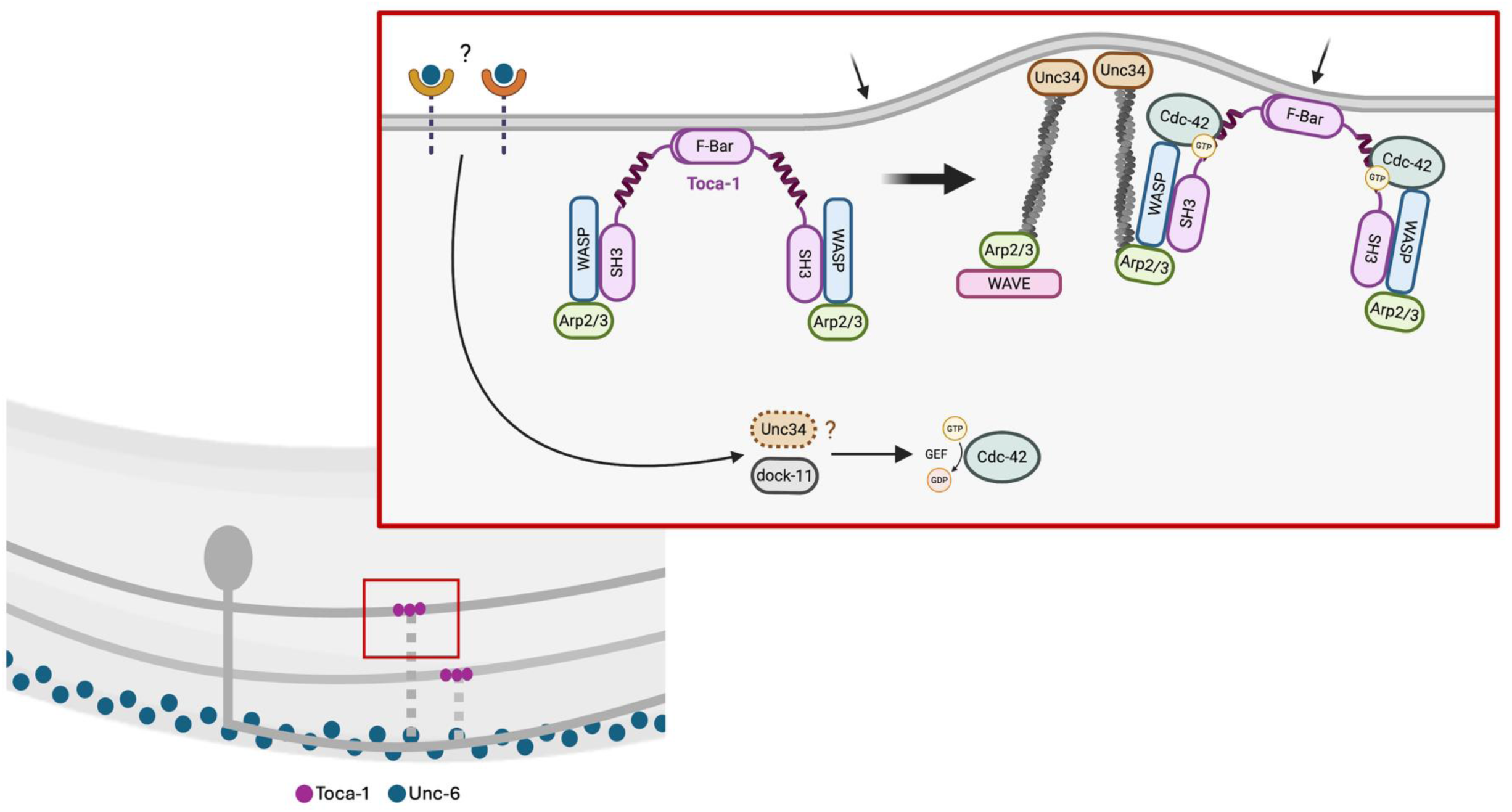
Model for TOCA-1-mediated PLM branch formation. TOCA-1 functions as an essential organizing protein that integrates membrane dynamics with actin regulatory pathways to control PLM collateral branching. DOCK-11 acts upstream as a CDC-42 GEF, potentially providing spatial regulation that determines where CDC-42 becomes activated along the PLM axon. TOCA-1 serves as a crucial organizing protein required for multiple downstream effectors: its F-BAR domain induces membrane curvature at branch initiation sites while its coiled-coil and SH3 domains recruit and organize CDC-42 and WSP-1 to drive localized Arp2/3-mediated actin nucleation. The model incorporates three distinct actin regulatory pathways that converge on branch formation: UNC-34/Ena/VASP functions in a parallel pathway to generate linear actin filaments for branch elongation and stabilization, while the WAVE complex provides additional Arp2/3 activation that acts redundantly with the WSP-1 pathway. Netrin signaling integrates with this intrinsic branching machinery to provide spatial information that influences TOCA-1 complex assembly or activity. Endocytic components (dynamin, clathrin) regulate the availability of membrane components, guidance receptors, or other nucleation factors, adding spatial and temporal control to promote branch formation at correct axonal positions. TOCA-1 may also be involved in this process, inducing membrane curvature and promoting endocytosis. Created using BioRender.

The model incorporates three distinct actin regulatory pathways that converge on branch formation. UNC-34/Ena/VASP functions in a parallel pathway to generate linear actin filaments necessary for branch elongation and stabilization (Bear *et al*., 2002; Withee *et al*., 2004), while the WAVE complex provides additional Arp2/3 activation that can act redundantly with the WSP-1 pathway (Shakir *et al*., 2008; Zhu *et al*., 2016). The integration of netrin signaling with this intrinsic branching machinery suggests that guidance cues may provide spatial information that influences TOCA-1 complex assembly or activity.

### Broader implications and future directions

Our work reveals fundamental principles of how neurons achieve precise spatial control over branching decisions. The identification of density-dependent assembly mechanisms for TOCA-1 regulation suggests that concentration-dependent thresholds may be a general strategy for restricting branching to stereotyped sites. Future work should examine whether similar mechanisms operate in other neuronal contexts and how spatial signals control the localization and activity of branching machinery.

Key outstanding questions include how DOCK-11 activity is spatially regulated to determine CDC-42 activation sites, how TOCA-1 localization is controlled, and how netrin signaling as well as endocytic mechanisms interface with the TOCA-1 pathway. Understanding these mechanisms will provide deeper insights into the spatial control of neuronal morphogenesis.

## Data Availability

Strains and plasmids are available upon request. The genotype of strains and sequence of alleles and plasmids are found in the Supplementary Table 1. The authors affirm that all data necessary for confirming the conclusions of the article are present within the article, figures, and tables.

## Acknowledgments

We thank Emma Knoebel technical assistant, Diana Shen for performing an initial version of *arp-2* auxin experiment, Barth Grant for providing the *pwSi540* Tir1 transgene and Chris Quinn for comments on the manuscript.

## Abbreviations

TRN: touch receptor neuron
AP: anterior-posterior
WASP: Wiskott-Aldrich syndrome protein
GEF: Guanine nucleotide exchange factor,
Arp: Actin related protein,
VC: ventral cord
RMCE: Recombination-Mediated Cassette Exchange
PCH: *pombe* Cdc15 homology
mNG: monomeric Neon Green FP
FP: Fluorescent Protein
VCA: Verprolin Central Acid
GFP: Green FP

